# Insulated *piggyBac* and FRT vectors for engineering transgenic homozygous and heterozygous eHAP cells

**DOI:** 10.1101/2024.04.19.590353

**Authors:** Annabel Y Minard, Stanley Winistorfer, Robert C Piper

## Abstract

Transgene expression in eHAP cells, a haploid cell line popularly used to generate gene knockouts, is difficult owing to its low transfection efficiency and propensity for silencing integrated transgenes. To simplify transgene expression, we engineered insulated integrating plasmids that sustain high levels of transgene expression in eHAP cells, and that can be used in other cell lines. These vectors are compatible with FLP-FRT and *piggyBac* integration, they flank a gene-of interest bilaterally with tandem cHS4 core insulators, and co-express nuclear-localized blue fluorescent protein for identification of high expressing cells. We further demonstrate that transgenic haploid eHAP cells can be fused to form transgenic heterozygous diploid cells. This method creates diploid cells carrying the transgenic material of the haploid progenitors and could also be used to create heterozygous cells of defined genotypes. These tools expand the repertoire of experiments that can be performed in eHAP cells and other cultured cells.

## Introduction

eHAP cells, a fully haploid version of HAP1 cells, are useful in gene-function studies because its haploid genome requires editing only one allele to obtain a gene deletion (1, 2). Most gene-function studies also require the expression of transgenes. This includes to complement a phenotype by re-expression of a gene-of-interest (GOI), or to monitor a phenotype with reporter constructs. During our studies, we noted conventional strategies achieve poor transgene expression in eHAP cells (3). Notably, the efficiency of transient transfection was low, and stably integrated transgenes underwent silencing. Here we overcome this limitation with vectors for simple and reliable transgene expression in eHAP cells. These vectors improve transgene expression by 1) using the *piggyBac* or Flp-In System to stably integrate transgenes, 2) incorporating tandem cHS4 core insulators that diminish epigenetic silencing, and 3) co-expressing nuclear-localized blue fluorescent protein to identify and sort for cells with transgene expression.

The *piggyBac* and Flp-In systems are non-viral methods of stable integration, which involve transfection of cells with a recombinase and expression plasmid. The expression plasmid contains a GOI, and DNA elements that stimulate the recombinase to integrate the GOI into the genome. With the Flp-In system, Flp-recombinase inserts a transgene into an Flp Recombinant Target (FRT) site in the host’s genome (4-6). The FRT site must be introduced prior to integration. Because integration occurs only at an FRT site, this method creates congenic stable cell lines. In the *piggyBac* system, *piggyBac* transposase integrates transgenes flanked by inverted terminal repeats (ITR) into sites throughout the genome, with a bias for active chromosomal regions rich in AATT sequences (7-9). Therefore, *piggyBac*-mediated integration does not produce congenic cells. So that cells with similar expression levels can be selected from a mixed population, we coupled the expression of the GOI with BFP via an Internal Ribosome Entry Site (IRES) in our vectors. By providing two strategies here, the user can choose the optimal strategy or use both to integrate two expression vectors.

Transgenes integrated by either system experienced epigenetic silencing in eHAP cells.

Epigenetic silencing is mediated by repressive covalent modifications on DNA and histones. These modifications promote the spreading of heterochromatin and repel transcription machinery (10-12). The causes of silencing are not always known, but can include the sequence, genomic position, and integration method of the transgene, as well as the cell type of the transgene host (13). One way to overcome gene silencing is by flanking a GOI with barrier insulators such as cHS4 (14). The majority of the insulating effect of the cHS4 insulator is contained within a 250 bp “core insulator” that recruits chromatin-modifying enzymes (15). Its potency is increased when present in tandem (16). We incorporated sets of tandem cHS4 core insulators into our expression-vectors and find this amplified stable gene expression.

Here we also demonstrate that transgenic haploid eHAP cells can be fused to create transgenic heterozygous diploid cells. This method allows for multiple integrated expression vectors, obtained from haploid transgenic progenitor cells, to be hosted in a diploid cell. It also allows for engineering of heterozygous cells with defined genotypes, which is useful for studying genetic interactions, particularly those that occur in compound heterozygous and polygenic diseases. In this method, isogenic haploid cells are fused via PEG 1500 and the resulting diploid cells can be selected for based on acquisition of antibiotic resistance and fluorescent protein markers from the haploid progenitors.

The methods presented here allow for engineering transgenic homozygous or heterozygous eHAP cells. The insulated expression vectors can also be used in other mammalian model systems that suffer gene silencing.

## Methods

### Cell culture

Human haploid eHAP cells (Cat # C669, Horizon Discovery, Cambridge MA) were cultured in Iscove’s Modified Dulbecco’s Medium (IMDM; Gibco, Billings, MT) supplemented with 10% FCS. Cells were passaged every 48-72 h. As noted previously, eHAP cells can diploidize (17). To enrich for haploid cells, we used flow-actuated cell sorting (FACS) using size (FSC: forward scatter; SSC: side scatter) as the sorting parameter. For antibiotic selection of stable transfectants, eHAP cells were grown in IMDM FCS with puromycin 500 µg/ml and/or neomycin 500 µg/ml. eHAP-FRT cells were generated as described previously (3). Briefly, eHAP cells were transduced with lentivirus carrying the pQCXIP FRT GFP-Neo^R^ (pPL6490) vector, which introduced the locus illustrated in Fig 1a.

**Fig 1.**
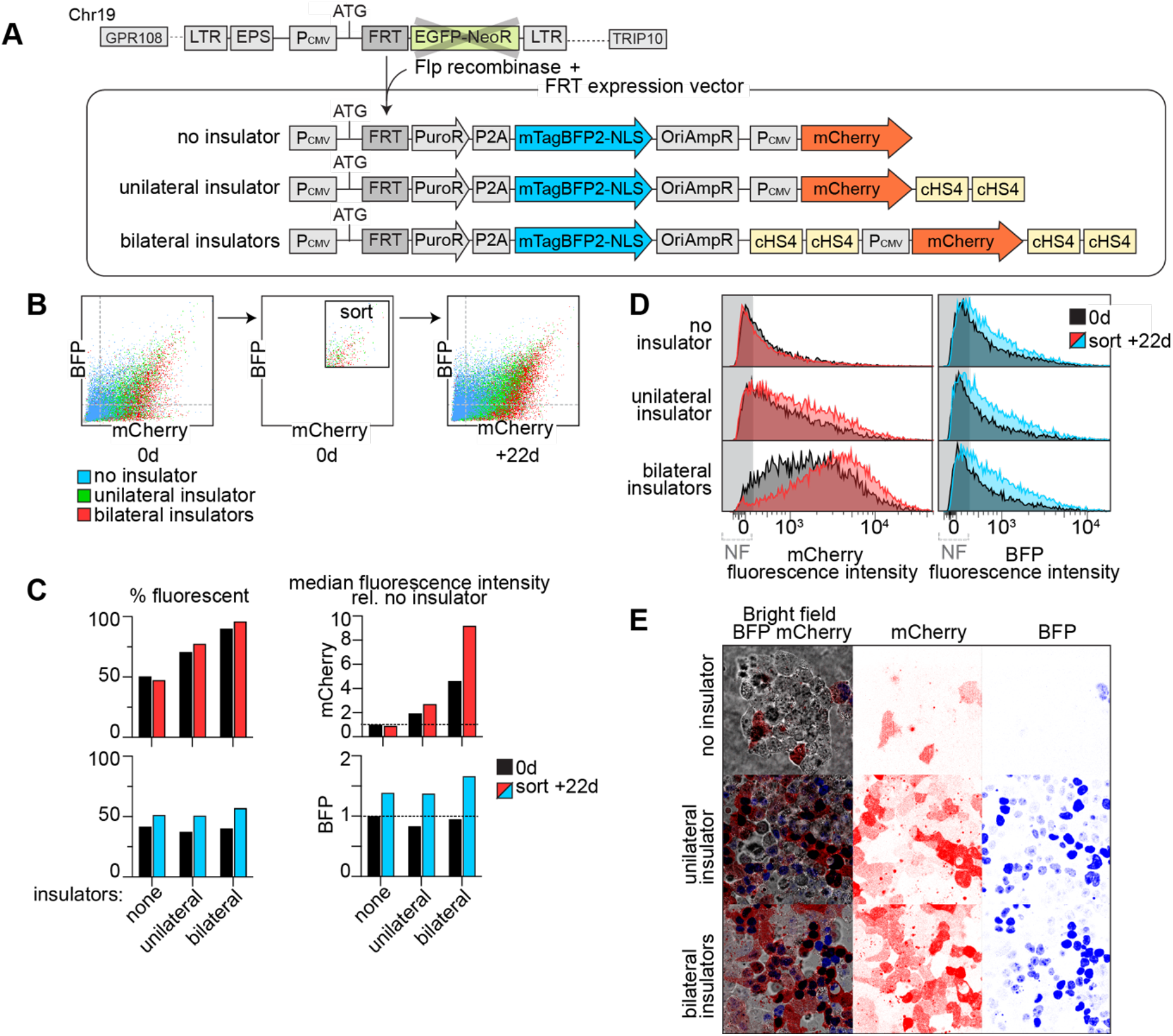
Insulated FRT vector rescues expression of GOI. **(A)** Schematic of FRT locus in eHAP FRT cells and the vectors used in this study. **(B)** After antibiotic selection of transgenic cells, the fluoresce intensity of mCherry and BFP were assayed by FACS. Bright cells were isolated by FACS, passaged for 22 d, and the fluorescence intensity was assayed again. **(C)** Quantification of the % fluorescent cells and median fluorescence of cells analyzed in (B). **(D)** Histogram of the distribution of fluorescence intensities of the cells assayed in (B). **(E)** Micrograph of transgenic cells after antibiotic selection.

### Plasmids

Flp-Recombinase Expression Vector pOG44 was from ThermoFisher. A hyperactive *piggyBac* transposase (18) in a pcDNA3.1 vector was used as described (19). The uninsulated FRT integration vector, single cHS4 FRT integration vector, double cHS4 FRT integration vector, uninsulated *piggyBac* vector, insulated *piggyBac* vector maps are attached in Supplemental materials. The vectors were sequenced using long-reading sequencing at plasmidsaurus (London, UK) and deposited in Addgene (Watertown, MA).

### Plasmid transfection

Transfection of eHAP cells was performed by reverse-transfection using Lipofectamine (20). Compared to standard-transfection, reverse-transfection achieved a higher efficiency while requiring half the amount of reagent and one less day of preparation. In our best protocol, the DNA-lipofectamine mixture was prepared by mixing 1µg DNA, 2 µl Lipofectamine 3000, 2 µl P3000 reagent, and 112 µl Opti-MEM, according to the manufacturer’s instructions (Invitrogen/Thermofisher, Waltham, MA). The DNA lipofectamine mixture was combined with 6 x 10^6^ trypsinized cells, 2 ml of IMDM and 10% FCS in a single well of a 6-well dish and incubated overnight.

Electroporations of eHAP cells were performed using the Lonza 4D Nucleofector. This system comes with three electroporation solutions and fifteen electroporation programs. Using the manufacturer’s Cell Line Optimization Kit, we determined that the best protocol combined the SF solution with EH-100 program. If using the other solutions, the SE solution paired best with DS-120 or E0-100 programs, and the SG solution paired best with EH-100 program.

### Fluorescence-activated cell sorting (FACS)

For cell sorting experiments, a Becton Dickinson Aria II or a Cytek Aurora equipped with a 130 µM diameter nozzle was used. FACS data was analysed using Flow Jo Version 10 (BD Biosciences) software.

### Karyotyping eHAP cells

The ploidy of eHAP cells was determined by propidium iodide staining. Cells were trypsinized, washed twice with PBS, lysed and stained using Nicoletti buffer (0.1% sodium citrate, 0.1% Triton X-100, 0.5 unit/mL RNase A, 20 units/mL RNase T1, 50 μg/mL propidium iodide). The intensity of propidium iodide stained was measured on a Becton Dickinson LSR II flow cytometer. Haploid eHAP cells were used for reference.

### Haploid Fusion

The protocol for cell fusion was adapted from the protocol by Yang, Shen (21). Haploid eHAP FRT-GFP cells and haploid PB tdTomato cells were grown to 50% confluency in a 10 cm^2^ dish. The cells were harvested by trypsinization, resuspended in IMDM FCS, pelleted at 100*g* for 5 min and resuspended in 5 ml IMDM without serum. 1 ml of resuspended cells from both cell lines were mixed, pelleted at 100*g* for 5min, resuspended in 50% PEG 1500 (Cat #10783641001, Roche, Indianapolis, In) and incubated for 2 min at room temperature. To this reaction, 10 ml IMDM was added, and incubated for 20 min at room temperature. Cells were pelleted at 100*g* for 5 min and resuspended in IMDM FCS and plated onto a 35 mM dish. The following day, cells were trypsinized and serially diluted to obtain single colonies in IMDM FCS with 500 ng/ml puromycin and 1 mg/ml neomycin. A colony expressing both GFP and tdTomato was isolated using disc cloning. The GFP and tdTomato expressing cell line was expanded. GFP and tdTomato expressing cells were enriched for by flow cytometry, expanded and karyotyped.

## Results

### Insulated FRT-expression vector for high levels of stable transgene expression

In our previous studies we optimized transient transfection protocols for eHAP cells, both by reverse-transfection with lipofectamine and by electroporation (see methods for protocols) (3). However, at best these methods achieved a transfection efficiency of 30%. Therefore, we developed systems for stable integration. First, we integrated an FRT site into eHAP cells to use the Flp-In system. As described below, genes integrated into the FRT locus experienced rapid gene silencing, and this necessitated the construction of vectors that insulate the GOI.

Previously, we introduced an FRT site into the eHAP genome by transducing eHAP cells with the pQCXIP FRT GFP-Neo vector by lentivirus (3). In this vector a CMV promoter drives expression of eGFP fused to a neomycin resistance gene (eGFP-Neo^R^). An FRT site is placed between the start codon and the eGFP-Neo^R^ ORF. Flp-recombinase-mediated integration of the pcDNA5 FRT expression vectors, as shown in Figure 1a, shifts the eGFP-NeoR out of frame, and in its place inserts an antibiotic resistance gene, in this case the hygromycin resistance gene (HygR). This is followed by a second CMV promoter that drives the expression of the gene of interest, in this case mCherry. Hygromycin resistant clones were isolated from multiple eHAP-FRT cell lines, however only two eHAP-FRT cell lines expressed mCherry and the level of expression was low. The brightest eHAP-FRT cell line was used in future experiments and the eGFP was deleted by CRIPSR to allow eGFP to be used in other constructs.

To ensure we were obtaining proper recombination of transgenes at the FRT locus, we replaced the hygromycin resistance gene in the integrating plasmid with the puromycin resistance gene fused to BFP with a nuclear localization signal (PuroR-BFP-NLS) (Fig. 1a). Expression of BFP-NLS indicates successful integration, since this ORF lacks a promoter and start codon and only expresses upon recombination at the genomic FRT locus. PuroR was used because it allows more rapid selection. Although we could observe BFP expression early after Flp-mediated integration, that fluorescence was lost upon passaging cells. This indicated that the FRT vector was integrated, but probably poorly expressed due to gene silencing.

To improve gene expression, we further modified the integrating vector with tandem cHS4 core insulators (16). The cHS4 insulator possesses barrier activity that blocks the spreading of heterochromatin and repressive epigenetic modifications (14). It also can block activity from nearby enhancers that might otherwise cause variable expression. We generated a vector in which tandem cHS4 core insulators were placed downstream of the mCherry cassette, and another vector in which tandem cHS4 core insulators were placed both upstream and downstream of the mCherry cassette (Fig. 1a). We compared the level of mCherry expression in the FRT integrating vectors by FACS (Fig. 1b). Without insulators, mCherry was expressed in only 51% of cells, the addition of one set of tandem cHS4 core insulators increased the mCherry expressing population to 71%, and with bilateral insulators the fluorescent population increased to 90% (Fig 1c-d). The median fluorescence of mCherry also increased as a result of the insulators, with unilateral insulators increasing fluorescence 1.9 fold, and bilateral insulators increasing fluorescence 4.6 fold. The expression of BFP-NLS was unaffected by the insulators, however it remained a useful indicator of successful integration and a predictor of mCherry abundance. We examined the fluorescence intensity of mCherry and BFP-NLS by microscopy (Fig. 1e). Without insulators the fluorescence of mCherry and BFP-NLS was barely detectable, in contrast, with the bilaterally insulated construct very bright cells were produced. Thus, the insulators improved gene expression, and bilaterally placed insulators surrounding the gene of interest were better than unilaterally placed insulators.

To obtain a population with an even higher level of expression, we enriched for the highest expressing mCherry and BFP cells by FACS (Fig. 1b). After passaging the sorted cells for 22 d, the bilaterally insulated construct produced a higher level of mCherry expression with 96% of cells expressing mCherry and a 9 fold increase in median fluorescence relative to the uninsulated construct at the start of the experiment (Fig. 1c-d). By comparison, the unilaterally insulated mCherry construct produced a population that was 78% fluorescent with a 2.7 fold increase in median fluorescence compared to the uninsulated construct at the start of the experiment. The uninsulated construct did not substantially change after sorting and passaging, 47% of cells were fluorescent and the median fluorescence intensity was 0.9 fold compared to the initial intensity.

In summary, the inclusion of bilateral tandem cHS4 core insulators improved gene expression substantially and was more beneficial that unilateral insulators. The bilaterally insulated construct increased fluorescence intensity 4.6 fold. With sorting the high level of mCherry expression is sustained for 22 d, and is increased 9 fold. The benefits of the insulators are restricted to the mCherry locus, and do not benefit the puroR-P2A-BFP-NLS locus. These results demonstrates that the bilaterally insulated FRT vector can be used to generate a high level of sustained expression of GOI.

### Insulated *piggybac* vector for selectable levels of stable transgene expression

We next sought to obtain high levels of stable gene expression using the *piggyBac* transposase system. We constructed a *piggyBac* expression vector that contained a CMV promoter driving the transcription of the GOI, in this case mCherry, followed by an IRES that coupled the levels of mCherry translation with PuroR-BFP-NLS (Fig. 2a). We coupled the expression of the GOI with BFP-NLS so that BFP fluorescence intensity could be used to account for variations in the expression of the GOI, or to select for populations with a desired level of expression. This feature is useful, since unlike the Flp-in system, *piggyBac* transposase integrates transgenes into AATT rich-sites throughout the genome, and each site may have individual affects on expression levels. We generated versions of this construct with and without insulators. The same insulators that were used in the FRT vectors were used here. The insulators were tandem cHS4 core insulators, placed both upstream and downstream of the bicistronic cassette (Fig. 2a). The vectors were co-transfected with a plasmid expressing *piggyBac* transposase into eHAP cells and a mixed population of puromycin resistant cells were selected for.

**Fig 2.**
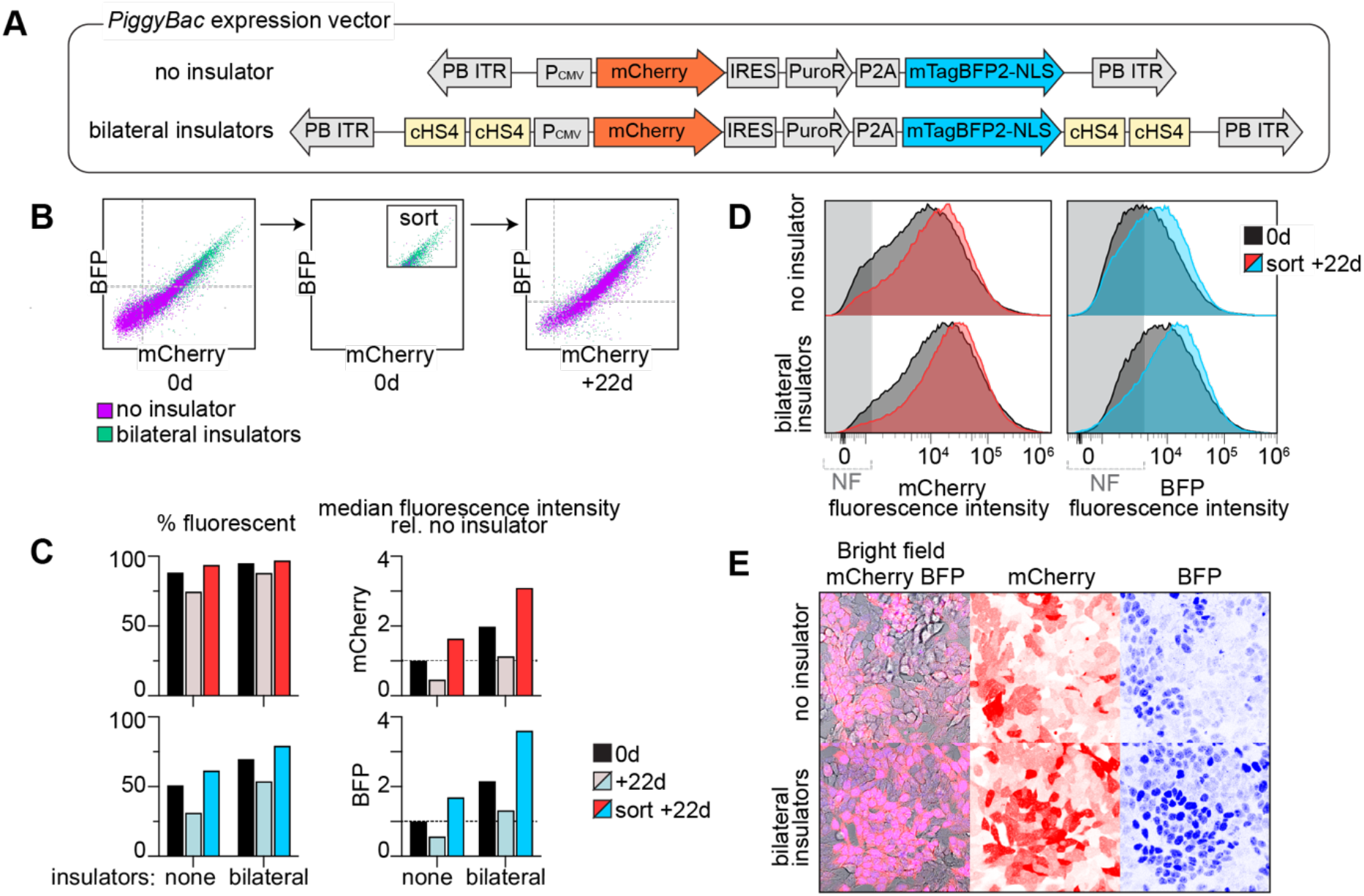
Insulated *piggybac* vector increases fluorescence of GOI. **(A)** Schematic of the *piggyBac* vectors used in this study. **(B)** After antibiotic selection of transgenic cells, the fluorescence intensity of mCherry and BFP were assayed by FACS. Bright cells were isolated, passaged for 22 d, and the fluorescence intensity was assayed again. **(C)** Quantification of the % fluorescent cells and median fluorescence of cells analyzed in (B). **(D)** Histogram of the distribution of fluorescence intensities of the cells assayed in (B). **(E)** Micrograph of transgenic cells after antibiotic selection.

The population of mCherry expressing cells in the uninsulated *piggyBac* vectors was higher than in the uninsulated FRT vectors, with 88% of cells expressing mCherry (Fig. 2b-e). This may be due to *piggyBac* vectors preferring to integrate at actively transcribed regions (7, 8). The insulated *piggyBac* vectors improved gene expression further, with 95% of cells expressing mCherry, and a 1.9 fold increase in mCherry median fluorescence intensity relative to the uninsulated construct. BFP expression correlated tightly with mCherry expression in both constructs, which allows it to be used to track the abundance of the GOI (Fig. 2b, e). By microscopy we also observed that compared to cells expressing the uninsulated construct, cells with the insulated construct had higher mCherry and BFP fluorescence (Fig. 2e).

To obtain a population with higher expression, we enriched for the highest expressing mCherry and BFP cells by FACS (Fig. 2b). We also examined if sorting improved the characteristics of the population by passaging unsorted cells for 22d. We found that enriching for high expressing cells was beneficial. In cells with the uninsulated constructs the median fluorescence increased 1.6 fold in the sorted group, compared to 0.45 fold drop in the unsorted group (Fig. 2c-d). The insulated vector produced a population with higher expression. In cells with the insulated constructs, the median fluorescence increased 3.1 fold in the sorted group, and 1.1 fold in the unsorted group.

Thus, the insulated *piggyBac* vector improves expression of the GOI, in this case mCherry, 1.9 fold, and with sorting 3 fold. BFP-NLS expression can be used to estimate mCherry expression levels.

### Engineering Heterozygous Diploids from Fusion of Transgenic Haploids

During our experiments, and as noted in other studies, we found that eHAP cells spontaneously diploidized and generate mixed aneuploid populations (17, 22, 23). We reasoned that directing diploidization in a controlled manner could provide an easy way to construct diploids of defined genotypes, such as heterozygous cells, or to introduce multiple GOIs into one cell line. In addition, we found that spontaneous diploids made from haploids, are larger and have different morphological characteristics that lend themselves better to certain experiments (3). These morphological differences were examined in detail by another group (17).

To accomplish directed diploidization, we constructed transgenic haploid cells lines with different drug selection markers and different fluorescent markers. For one haploid cell line we used the haploid eHAP-FRT cells expressing eGFP-NeoR. For another, we used an eHAP cell line expressing Td-Tomato and puromycin resistance using the *piggyBac* system described above. We fused the cell lines using a method based on polyethylene glycol (PEG), which has been used in other cell types (21). After incubation with PEG, we selected for fused hybrid cells using both neomycin and puromycin (Fig. 3a). After isolating NeoR+/PuroR+ clonal cell lines, we performed flow cytometry analysis to verify which clones had acquired both GFP and tdTomato from the haploid progenitors (Fig. 3b) and were diploid (Fig. 3c). Figure 3 demonstrates the production of a hybrid diploid, with the predicted diploid DNA content and expression of GFP and tdTomato.

**Fig 3.**
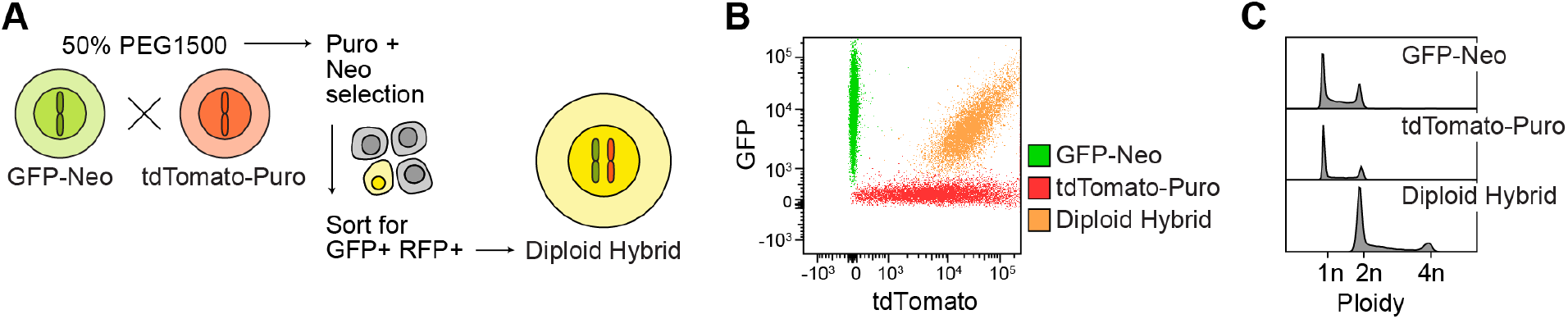
Heterozygous diploids engineered from haploid fusion. **(A)** Haploid eGFP-FRT-Neo cells and haploid PB-tdTomato-Puro cells were fused with 50% PEG1500. Fused cells were selected for in puromycin and neomycin, isolated by disc cloning and enriched by flow cytometry. **(B)** The fused cells expressed GFP and tdTomato as assessed by flow cytometry, **(C)** and were diploid as indicated by DNA content. DNA content was measured by quantifying PI staining intensity using flow cytometry.

## Discussion

Here we describe vectors that enable high levels of stable transgene expression in eHAP cells and that can also be used in other cells. These vectors stably integrate transgenes via the site-selective Flp-recombinase or *piggyBac* transposase. Expression of the transgene is sustained by cHS4 insulators and can be monitored by a co-expressed nuclear-localized BFP. The vectors assemble components that have been described earlier to create an effective gene delivery system in eHAP cells. These vectors may also be helpful in other cells lines with a high propensity for gene silencing.

These vectors were necessary because eHAP cells exhibited a high degree of silencing. The factors which contribute to silencing are diverse and not yet fully understood. Among the factors proposed to contribute to silencing are the sequence (e.g. presence of CpG islands), genomic position (e.g. which influences susceptibility to encroachment of heterochromatin), integration method of the transgene (which could activate viral and transposon defense systems), proliferation rate (which could cause antagonism between replication and transcription machinery), and cell type (they have varying levels of gene silencing machinery) (13). In our experiments the FRT integration vector experienced more severe silencing than the *piggyBac* integration vector. This may be because the FRT integration site was introduced by lentivirus and lentiviral sequences are well-documented to activate silencing (24, 25). It could also be that *piggyBac* integrates in more actively transcribed regions of the genome (7, 8). We found that cHS4 insulators were effective in eHAP cells, but they are not universally useful for overcoming gene silencing. In some cell lines cHS4 insulators in *piggyBac* vectors were reported to be beneficial (26, 27), while in others no benefit was observed (28). Thus, methods for transgene expression need to be optimized for each model system.

We also demonstrate that haploid eHAP cells can be fused to produce heterozygous diploid cell lines with defined genotypes. Haploid fusion is a potentially useful technique. It provides a means for introducing multiple expression vectors in one host, which is useful since the piggyBac and Flp-In system only support integration of one expression vector each per cell line. It can also be used to carry out haploinsufficiency screens, as demonstrated in a recent study using haploid embryonic stem cells (29). In that study, HVJ envelope particles were used to stimulate fusion. Fusion of haploid cells with different edited genes is also an easy method to model compound heterozygote and polygenic disease associated diseases variants. Engineering heterozygous cells using diploid cells is problematic due to the loss of heterozygosity that occurs during CRISPR gene editing (30). New methods to overcome this limitation have been developed. The new methods involve simultaneous delivery of HDR templates for both alleles (31, 32). Editing haploids, however, is potentially advantageous since only one allele is edited per cell. And when comparing multiple allele combinations, alleles do not need to be rederived, which also allows for a more consistent genomic background.

In conclusion we have presented methods for effectively generating transgenic eHAP cells and for making heterozygous diploid eHAP cells. The methods here expand the experiments that are possible to conduct in eHAP cells.

## Acknowledgement

We would like to thank John Lueck for advice on *piggyBac* and the University of Iowa Flow cytometry core which is supported by the Carver College of Medicine and the instrument grant 1S10OD034193-01. This work was supported by NIH RO1GM058202 to RCP. AYM was supported by ADA postdoctoral fellowship.

